# Changes in TP53 gene, telomere length, mitochondrial DNA, and its amount in benign prostatic hyperplasia patients

**DOI:** 10.1101/2024.05.07.592713

**Authors:** Egija Zole, Edgars Baumanis, Lauma Freimane, Rolands Dāle, Andrejs Leiše, Vilnis Lietuvietis, Renāte Ranka

## Abstract

Benign prostatic hyperplasia (BPH) is a growing issue due to an ageing population; however, there are not enough studies connecting it with ageing hallmarks. Some of the ageing hallmarks are changes in telomere length (TL) and genetic changes like possible mutations in the *TP53* gene or mitochondrial genome (mtDNA). Our study investigated these factors to see if they could be linked with BPH.

Prostate tissue samples were obtained from 32 patients with BPH (and 30 blood samples). As a healthy control group, age-matching blood DNA samples were used. For mtDNA sequence data comparison, 50 samples of a general Latvian population were used. The full mtDNA genome was sequenced using Next Generation Sequencing (NGS), for *TP53* gene – Sanger sequencing, and for mtDNA amount and telomere length – qPCR assay.

BPH patients in prostate tissue had much higher TL than in blood cells, and the healthy controls had the shortest telomeres. Also, the mtDNA amount in BPH prostate tissue was the highest compared with blood, and controls had the smallest amount. In the *TP53* gene, we did not find any mutations that could be linked to BPH. In the mtDNA genome, we found several unique mutations and heteroplasmic changes, as well as changes that have been linked before with prostate cancer (PC).

In conclusion, prolonged telomeres and changes in mtDNA amount might be involved in the development of BPH. Concerning mtDNA genome changes, some of the heteroplasmic or homoplasmic variants might contribute to the development of BPH. More studies should be conducted to prove or disapprove of these connections.

## Introduction

Benign prostatic hyperplasia (BPH) or benign prostate gland enlargement is due to unregulated hyperplastic growth of the epithelial and fibromuscular tissues and is an age-related disease. BPH is characterized by a proliferation of the prostatic stromal and epithelial cells; thereby, the BPH nodules increase in size, hence the difficulty of urinating. BHP can be observed in the vast majority of men as they age, especially those over 70. The development of BHP also depends on lifestyle, genetics, inflammation, and even geography. As the world is getting increasingly older, BPH becomes an increasing problem (reviewed in Chughtai et al., 2016).

However, there are still a lot of unknowns about the mechanisms of progression and biomarkers to predict the clinically significant disease. As it is an age-related disease, many cell-ageing hallmarks, like telomere shortening and mitochondrial dysfunction, could be involved in its development. It is well known that critically shortened telomeres trigger double-strand breaks signalling cascade and genomic DNA instability, suppress PGC-1α/β (peroxisome proliferator-activated receptor gamma, coactivator 1 alpha, and beta) action via the p53 transcription factor, causing increased p53 activity and high levels of apoptosis in which mitochondria are (Chin et al., 1999; Flores and Blasco, 2009; Sahin et al., 2011; Sahin and DePinho, 2010; reviewed in Zole and Ranka, 2018). *TP53* mutations can also influence mtDNA (mitochondrial DNA) maintenance, for example, *TP53*-R175H mutation could maintain mtDNA integrity, while *TP53*-C135Y induces greater mtDNA instability (Ericson et al., 2012; Zhuang et al., 2013). Mutations in the *TP53* gene can influence not only mtDNA maintenance but also mitochondrial biogenesis and function, and cell metabolism (reviewed in Giorgi et al., 2016; Kamp et al., 2016). However, interaction among these mechanisms is not widely studied in BPH.

mtDNA copy numbers or the amount has been studied in different diseases, cancer, and ageing, including BPH (reviewd in El-Hattab and Scaglia, 2013; Hu et al., 2016; Zole et al., 2018). Previously it has been shown that BPH patients have no different levels of cell-free mtDNA in plasma compared to healthy controls (Ellinger et al., 2008). In another study, BPH patients’ blood had slightly less mtDNA amount than prostate cancer (PC) patients, and cancer patients with higher Gleason scores were more likely to have higher mtDNA amounts than those with lower (Abhishek et al., 2017). Mutations and haplogroups of mtDNA have not been studied much in BPH; only one study was conducted with African men with or without BPH or PC (McCrow et al., 2016).

Another genome structure that has been involved in prostate dysfunction is the dysfunction of telomeres, which is generally associated with DNA damage and ageing and may cause chromosomal instability. There are controversial results in studies about telomere length (TL) in BPH patients. BPH typically is not associated with telomerase activity (an enzyme that prolongs telomeres) (Sommerfeld et al., 1996; Zhang et al. 1998; Mahjoub and Krams 2006), although in several studies some activity of telomerase or hTR and hTERT, expression has been shown in BPH tissue (Kamradt et al., 2003; Rane et al., 2016; Wymenga et al., 2000). In contrast, PC is strongly associated with telomerase activity (Wymenga et al., 2000; Bettendorf et al. 2003) but telomere shortening (Sommerfeld et al., 1996; Koeneman et al. 1998). In two studies, BPH patients had shorter telomeres in comparison to patients with normal prostate. In PC tissues, telomeres were the shortest (Cheng et al., 2020; Sommerfeld et al., 1996). In another study, it was shown that BHP patients had the longest telomeres in comparison to normal tissue and cancer tissues. Patients with BPH plus PC had shorter telomeres than patients only with BHP. The shortest telomeres were in PC tissues (Heaphy et al., 2010).

*TP53* mutations are reported in both PC (Ecke et al., 2010; Isaacs et al., 1991) and BPH (Meyers et al., 1993; Wertz et al., 1996), which might be a tumour risk factor (Schlechte et al., 1998). Mutated *TP53* can also affect TL, like a mutant *TP53*-R175H showed longer telomeres than the mutant *TP53*-V143A, which had longer than WT (wild type) in LoVo cell lines (Samassekou et al., 2014) but not in every case *TP53* mutations affect TL (Rampazzo et al., 2010).

So far, there has not been an article that covers the dynamics of telomeres and mtDNA, and the *TP53* gene together in one sample cohort. In our study, we investigated correlation and changes in TL and mtDNA amount, including unique variations in *TP53* and mtDNA in our BPH sample cohort.

## Materials and methods

### Sample collection, storage, and DNA extraction

Of the 32 prostate tissue samples in our study, 30 matched blood samples were available. All the tissue samples were obtained from patients with BPH during routine prostate operations; patients’ age from 59 to 85 years old, average age 72 years old. Prostate-Specific Antigen (PSA) levels were from 0.818 to 62.47 ng/ml. BPH diagnosis was confirmed, and cancer excluded by an anatomical pathologist. Informed consent was obtained from all patients, and the health history information was collected via the questionnaire. One PC tissue sample was used as a reference for qPCR telomere and mtDNA amount analysis. This sample was obtained from one of the BPH patients who, after the biopsy, was diagnosed with PC.

DNA samples from blood from healthy age-matching males were used as a control group (n=47, 59-85 years old). These samples were from individuals without prostate diseases, any cancer or other diagnosed telomere or mitochondrial disorders and were obtained from the Genome Database of the Latvian Population (VIGDB, bmc.biomed.lu.lv/lv/par-mums/saistitas-organizacijas/vigdb/).

To compare the BPH patents’ mtDNA genome with the general Latvian population, Next Generation Sequencing (NGS) mtDNA sequencing data from 50 individuals were used. The sequence data was obtained from VIDGDB; the average age was 48 years; 23.5% females (average age 53 years), 75.5% males (average age 47 years); samples without cancer or prostate diseases at the sampling time.

All data about the participants were kept fully anonymous. The study protocol was approved by the Central Medical Ethics Committee of Latvia, Nr.01-29.1/3. All tissue and blood samples were stored at −70 °C.

The total genomic DNA was extracted simultaneously from the prostate tissue and blood cells. Prostate tissue samples were prepared for DNA extraction by chopping the tissue into pulp. The pulp was placed into a tube with 2.5 mL of Tissue lysis buffer (10 mM Tris-HCl (pH=8), 10 mM EDTA (pH=8), 100 mM NaCl, 0.5% SDS). Then 25 µl of Proteinase K (20 mg/ml) was added and mixed. The tubes were incubated for 24h at +50 °C. Blood samples were prepared for DNA extraction by centrifuging vacutainers at 2500rpm/20min, at +4 °C. A plasma layer was removed, the rest of the blood was transferred to a 15 ml falcon, and red blood cells were removed by using Red blood cell lysis buffer A1 (0.32 M saccharose, 10 mM Tris-HCl, pH=7.6, 5mM MgCl_2_, 1% Triton X-100). Leucocytes were resuspended in 2.5 mL Cell Suspension solution (25 mM EDTA, pH=8.0, 75 mM NaCl) and incubated for 5 min at room temperature. Then 250 µL of 10% SDS and 4.5 µL of Proteinase K (20 mg/ml) were added, and the mixture was incubated for 50 min at +50 °C. DNA was extracted using the standard phenol-chloroform method as previously described (Sambrook et al. 1989). The DNA samples were stored in TE buffer (10 mM Tris–HCl, 1 mM EDTA, pH=8.0) at −20 °C. Absorbance readings (260 nm) of DNA extracts indicated the DNA concentration to be in the range 150–600 ng/µl.

### Relative qPCR SYBR green telomere length quantification assay

The ΔC_T_ method using a reference gene was used to measure TL in samples. qPCR was performed using Maxima SYBR green qPCR Master Mix (2X) (Thermo Scientific, USA). Telomeres-specific forward and reverse primers for one reaction were as follows: Telo1 (200 nM), 5′-GGTTTTTGAGGGTGAGGGTGAGGGTGAGGGTGAGGGT-3′, and Telo2 (200 nM), 5′-CCCGACTATCCCTATCCCTATCCCTATCCCTATCCCTA-3′. The quantitative PCR included an initial denaturation for 10 min at 95 °C, followed by 40 cycles at 95 °C for 10 s, and 58 °C for 1 min. TL measurements were normalized using the following β-globin gene-specific forward and reverse primers in a separate qPCR run: Beta-glob1 (300 nM), 5′-GCTTCTGACACAACTGTGTTCACTAGC-3′, and Beta-glob2 (500 nM), 5′-CACCAACTTCATCCACGTTCACC-3′. For qPCR, after a denaturation step at 95 °C for 10 min, samples were incubated for 40 cycles at 95 °C for 10 s and at 56 °C for 20 s (Kim et al., 2013). The concentration of the DNA samples for all qPCR reactions was 10 ng/µl in a 10 µl reaction mixture. Each sample was run in triplicate. A no-template control and duplicate calibrator samples were used in all runs to allow comparisons of the results across all the runs. A melting curve analysis was performed to verify the specificity and identity of the PCR products. TL was calculated using threshold cycle or C_T_ values and the following equation: relative TL ratio_(test/reference)_=2^Ct(β-globin)−Ct(telomeres)^.

### Relative qPCR TaqMan mtDNA copy number quantification assay

Relative mtDNA copy number was measured by using qPCR with the Maxima Probe/ROX qPCR Master Mix (2X) (Thermo Scientific, USA). MtDNA copy number amount was normalized by simultaneous measurements of the nuclear gene *GAPDH* and the mitochondrial D-loop products. The forward and reverse primers (1250 nM each) for the *GAPDH* reaction were GapdhF, 5′-GAAGGTGAAGGTCGGAGT-3′, and GapdhR, 5′-GAAGATGGTGATGGGATTTC-3′, respectively; the TaqMan probe (250 nM) was GapdhTqM, 5′-CAAGCTTCCCGTTCTCAGCC-3′. The forward and reverse primers (50 nM each) for the mitochondrial D-loop were FmtMinArc, 5′-CTAAATAGCCCACACGTTCCC-3′, and RmtMinArc, 5′-AGAGCTCCCGTGAG TGGTTA-3′, respectively; the TaqMan probe was PmtMinArc (250 nM) - 5′-CATCACGATGGATCACAGGT-3′ (Phillips et al., 2014). The DNA concentration was 10 ng/µl in a 15 µl reaction. After a denaturation step at 95 °C for 10 min, the DNA samples were incubated for 40 cycles at 95 °C for 15 s, 57 °C for 30 s, and 72 °C for 30 s. Each sample was run in triplicate. A no-template control and duplicate calibrator samples were used in all runs to allow for the comparison of results across the runs. MtDNA copy number was calculated using threshold cycle values and the following equation: relative copy number ratio_(test/ref)_=2^Ct(Gapdh)−Ct(D-loop)^.

### PCR amplification and sequencing of exons 1–11 of the *TP53* gene

Primers used to amplify exons 1–11 of the *TP53* gene were described previously (Liu and Bodmer, 2006). PCR was performed in a final volume of 12.5 µl. The reaction mixture contained 1x Reaction buffer BD (0.8 M Tris-HCl, 0.2 M (NH_4_)_2_SO_4_), 2.5 mM MgCl_2_, 0.1 mM dNTPs mix, 400 mM of each primer, 1.25 U of FIREPol DNA Polymerase (Solis BioDyne, Estonia) and 20 ng of genomic DNA. The PCR assays were performed under the following conditions: initial denaturation at 95 °C for 5 min; 30 cycles of denaturation at 95 °C for 30 s, primer annealing at 55-60 °C for 30 s, and elongation at 72 °C for 38 s; and a final elongation step at 72 °C for 5 min. PCR products were separated using electrophoresis on a 1 % agarose gel (TopVision Agarose, Thermo Scientific, USA) in Tris–Acetate-EDTA buffer containing 0.2 µg of ethidium bromide/ml and were visualized with transillumination under UV light. Samples were enzymatically cleaned using Exonuclease I and FastAP Thermosensitive Alkaline Phosphatase (Thermo Scientific, USA). The sequencing reaction was performed by using ABI PRISM BigDye Terminator v3.1 Ready Reaction Cycle Sequencing Kit (Thermo Scientific, USA) and the same set of primers in 25 cycles under the following conditions: 94 ◦C for 30 s, 55 ◦C for 15 s, and 60 ◦C for 4 min. The sequenced material was analyzed by a standard technique using an ABI Prism 3100 Genetic Analyzer (Perkin-Elmer, USA). The BLAST program (http://www.ncbi.nlm.nih.gov/BLAST) was used for the comparison of sequences obtained in this study versus those previously deposited in GenBank.

### Full mtDNA genome sequencing by Next Generation Sequencing

The full-length mtDNA genome was analyzed by amplification of mtDNA using Thermo Scientifc Phusion High-Fidelity DNA Polymerase protocol (Thermo Scientifc, USA). Pre-amplification was applied to eliminate mtDNA-derived pseudogenes in the nuclear genome (NuMTs). PCR primers were described previously in Fendt et al.: FampA, AAATCTTACCCCGCCTGTTT; RampA, AATTAGGCTGTGGGTGGTTG; FampB, GCCATACTAGTCTTTGCCGC; RampB, GGCAGGTCAATTTCACTGGT (Fendt et al., 2009). PCR was performed in a final volume of 13 µl. The reaction mixture contained 1 x GC buffer, 200 µM dNTPs mix, 0.5 µM of each primer, 1 U of Phusion High-Fidelity DNA Polymerase and 20 ng of genomic DNA. The PCR assays were performed under the following conditions: initial denaturation at 98 °C for 30 s; 30 cycles of denaturation at 98 °C for 10 s, elongation 72 °C for 8 min, 15 s; and a final elongation step at 72 °C for 10 min. PCR products were separated using electrophoresis on a 1 % agarose gel (TopVision Agarose, Thermo Scientific, USA) in Tris–Acetate-EDTA buffer containing 0.2 µg of ethidium bromide/ml and were visualized with transillumination under UV light. Positive amplicons were sheared by sonication and prepared for Next Generation Sequencing (NGS) using Ion Xpress™ Plus Fragment Library Kit (Thermo Scientifc, USA) and NucleoMag® NGS Clean-up and Size Select beads (Macherey-Nagel, Germany) following manufacturers’ instructions. The 200 bp long libraries were barcoded (Ion Xpress^TM^ Barcode Adapters Kits) and sequenced on the Ion Personal Genome Machine (PGM^TM^) system using the Ion 314^TM^ Chip. For each 32 blood and 30 prostate tissue samples at least 83.2 Mb of data were generated with an average coverage depth of ∼ 5,000× per sample.

### NGS data analysis

To analyze NGS data usegalaxy.org (Afgan et al., 2018) and Integrative Genomics Viewer (IGV) (Robinson et al., 2011) were used. For mtDNA haplogroups and SNPs (single nucleotide polymorphisms) analysis HaploGrep 2.0 (Kloss-Brandstätter et al., 2011) and PolyTree.org (Van Oven and Kayser, 2009) were used. The sequences were analyzed by comparison with the revised Cambridge Reference Sequence (Andrews et al., 1999).

### Statistical analysis

A two-sided Chi-square test was used for nominal data statistics. After frequency distribution was tested, data did not show normal distribution and Mann– Whitney U test was used, and t-test was used by GraphPad Prism version 5 for Windows (La Jolla California USA, www.graphpad.com 2016). Data were expressed as means ±SEM (Standard Error of the Mean), and differences of *P*<0.05 were considered significant.

## Results

### Telomere length in blood and tissue samples of benign prostatic hyperplasia

To test if there is a difference in TL among healthy controls and BPH patients, TL was measured by qPCR in all the BPH prostate tissue, BPH blood, control group blood and the reference PC prostate tissue sample (**Figure 1**). A significant difference was observed between TL measured in blood cells of BPH patients and the healthy control group (P=0.0006), where BPH patients had longer telomeres (mean: 0.06713 ±0.1447 SD, median: 0.0325, range 0.015-0.055 ru (relative units)) than the control group (mean: 0.02634 ±0.007836 SD, median: 0.0250, range 0.018-0.785 ru).

**Fig. 1.**
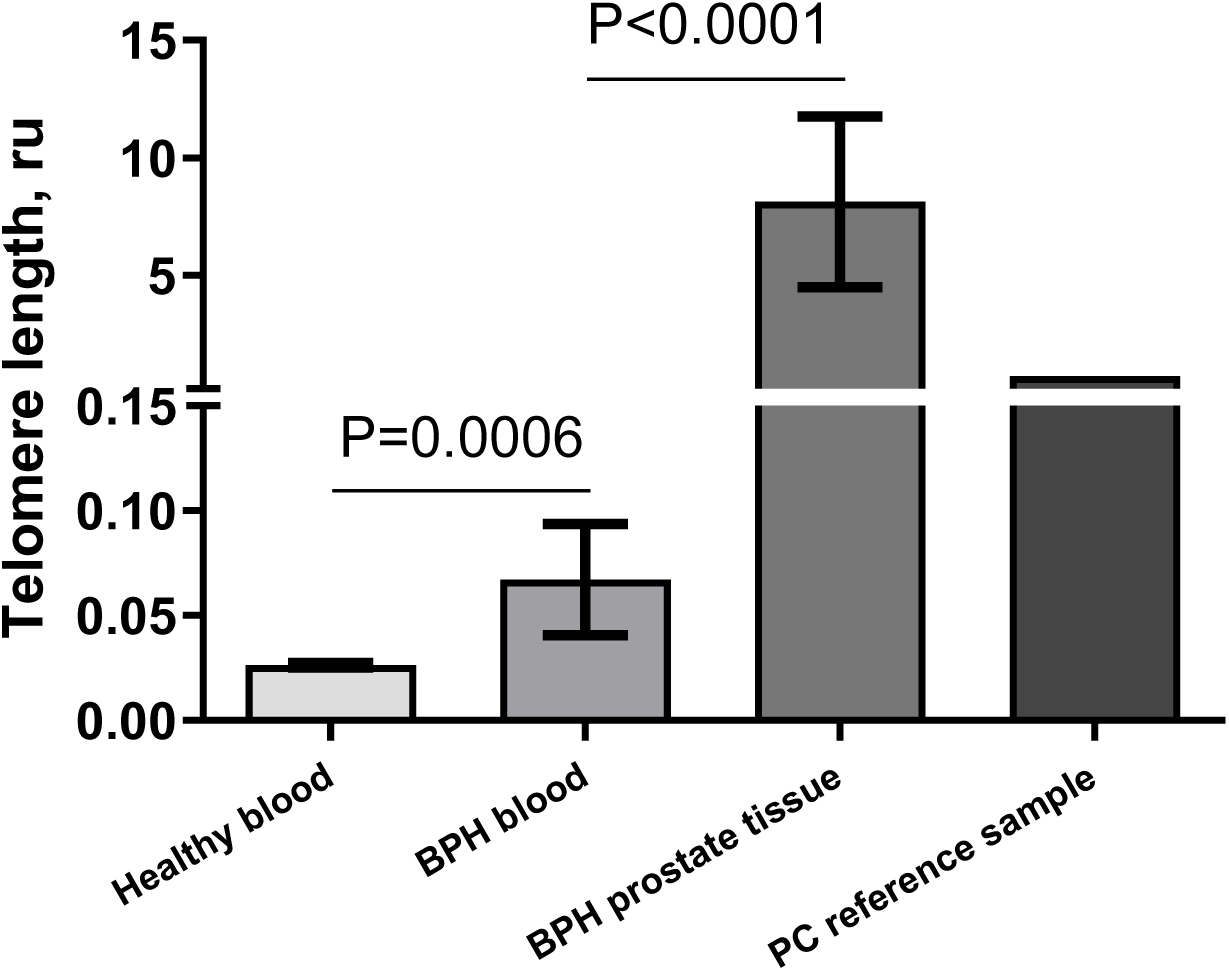
Telomere length in benign prostatic hyperplasia samples. BPH - benign prostatic hyperplasia, ru - relative units, data were expressed as mean ±SEM

For BPH patients, TL was significantly longer in prostate tissue (mean: 8.126 ±20.56 SD, median: 0.6825, range 0.046-83.58 ru) than blood, P<0.0001, while TL in the PC tissue reference sample was 0.704 ru. TL was more heterogeneous in BPH patients’ prostate and blood samples than in the controls’ blood samples. There was no connection between TL and PSA levels in our sample cohort. These results confirms that there are changes in TL in BPH disorder.

### Mutations in *TP53* gene in benign prostatic hyperplasia patients

To find mutations in the *TP53* gene, all 11 exons located on chromosome 17p13.1 were sequenced by Sanger sequencing for all the BPH blood and prostate tissue and healthy control group blood samples. A missense mutation at position 7676154C>G in exon 4 was the only mutation found. It changes 72^nd^ amino acid of the TP53 protein from a proline (cyclic amino acid) to an arginine (basic amino acid). There was no difference between BPH blood and tissue samples. Among BPH patients, only 1 (3.1%) was homozygous for wild-type TP53 gene sequence (WT), while a heterozygotic variant was detected in 14 (43.8%) samples, and 17 (53.1%) patients were homozygous for 7676154C>G allele.

Also, there was a mutation in an intron between exons 2 and 3 (7676483C>G) that represented the same heterozygotic and homozygotic findings as the 7676154C>G mutation; thus, the mutation could be considered as an inherited and a frequent SNP in a Latvian population. To test this hypothesis, the *TP53* gene was sequenced in all the healthy controls’ (n=47) DNA samples; 4 (8.5%) had homozygotic WT variant, 19 (40.4%) had the heterozygotic variant, and 24 (51.1%) individuals had the homozygotic mutated variant, showing no connection with BPH development. There was no difference in TL, mtDNA amount or PSA level between *TP53* heterozygotic or homozygotic samples (data not shown).

### Mitochondrial DNA amount in blood and tissue samples of benign prostatic hyperplasia

To test if there is a difference in mtDNA amount, it was measured by qPCR in all BPH prostate tissue, BPH blood cells, control group blood cells and the PC prostate tissue samples (**Figure 2**). On average, BPH patients’ blood cells had more mtDNA (mean: 39.96 ±16.50 SD, median: 38.37, range 14.40-72.62 ru) than the control group (mean: 28.67 ±11.13 SD, median: 26.63, range 14.70-70.44 ru), P=0.003. Prostate tissue samples had more mtDNA (mean: 168.7 ±63.40 SD, median: 160.6, range: 60.87-425.69 ru) than patients’ blood cells, P<0.0001. The PC reference sample (204.91 ru) had more mtDNA than BPH prostate tissue samples. There was no connection between mtDNA amount and PSA levels in our sample cohort. These results show that there are changes in mtDNA copy number in BPH disorder.

**Fig. 2.**
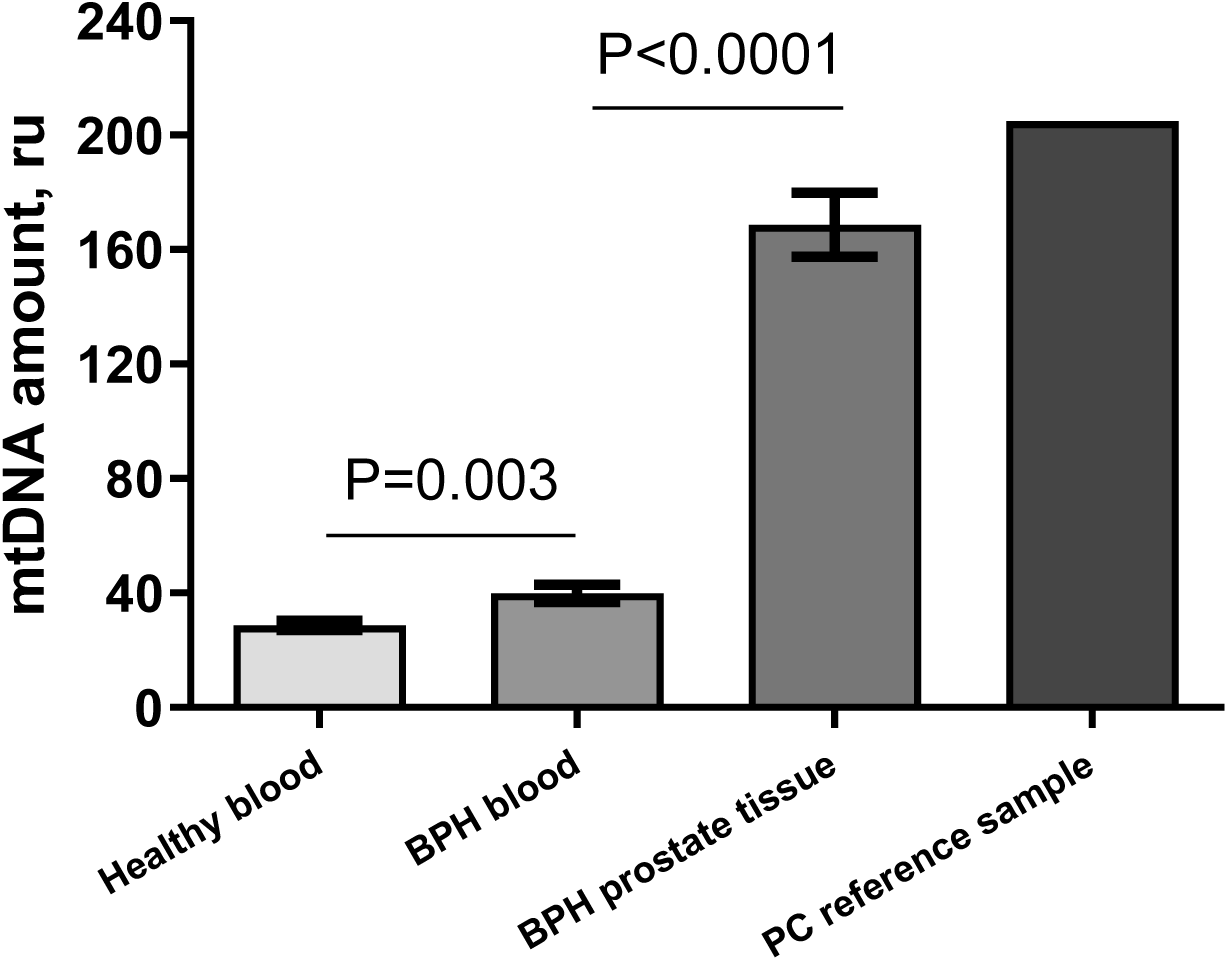
Mitochondrial DNA (mtDNA) amount in blood and prostate tissue samples of benign prostatic hyperplasia patients. BPH - benign prostatic hyperplasia, ru – relative units, data were expressed as mean ±SEM

### Full-length mtDNA analysis in benign prostatic hyperplasia

To analyze mutation and haplogroups of mtDNA, the full length of mtDNA was sequenced by using the NGS method for all the BPH prostate tissue samples and blood samples. In BPH group MtDNA haplogroup U were 34% in comparison to previously reported in literature in the Latvian population 25% (Zole et al., 2015) (P=0.346, Chi-square=0.8890, df=1, OR=1.430). The rest of the haplogroups detected in the patients: H-34%, HV-6%, I-6%, J-6%, N-3%, T-9%. Overall, these results are similar to those obtained for the Latvian population.

For heteroplasmy level, the 5% threshold was set during the bioinformatics analysis in usegalaxy.com, i.e., the heteroplasmy level was considered significant if the mutation was present in ≥5% of reads, as the low heteroplasmy level might not have an effect on functions of the cell (Saneto, 2017), and also to avoid possible sequencing data errors and NuMTs. If a heteroplasmy was detected above 5% threshold level, then for those SNPs, the threshold was removed to get a more precise analysis.

In this study, several homoplasmic and heteroplasmic mtDNA mutations were detected in BPH patients’ samples. After comparing the sequence data of the 50 samples of the general Latvian population and the available literature about mtDNA variants in PC and BPH patients, all detected mutations were divided into 3 groups. The first group of 2 mutations from our BPH samples were heteroplasmic variants that were not detected in any of the samples from the general Latvian population and have been mentioned previously in the literature associated with PC (reviewed in Kalsbeek et al., 2017) plus are not associated with any of a mitochondrial haplogroup by PhyloTree.org (**Table 1**). The mutations were in 16S and tRNA-Ala regions.

**Table 1.** mtDNA variants, previously associated with PC that were detected in our sample cohort.

| POS | Allele change | Sample count that had the mutation (occurrences) | Heteroplasmy (cases) |  | Gene | Amino acid change | Article | Prediction in the publication |
| --- | --- | --- | --- | --- | --- | --- | --- | --- |
|  |  |  | Blood | Prostate tissue |  |  |  |  |
| 3047 | G>A | 5 | no<br>1%<br>no<br>1%<br>3% | 6%<br>no<br>1%<br>no<br>no | 16S | rRNA | Ju et al. 2014 | primary PC |
| 5590 | G>A | 4 | no<br>no<br>23%<br>no | 1%<br>1%<br>1%<br>1% | tRNA-Ala | acceptor stem | Ju et al. 2014 | primary PC |

\*These SNPs are not associated with any of a mitochondrial haplogroup by PhyloTree.org.
\*\*If heteroplasmy level for a SNP was detected above 5% threshold, then for those SNPs the threshold was removed to get more precise analysis.

The second group (**Table 2**) shows 30 homoplasmic or heteroplasmic mutations (together 86 individual occurrences of the 30 mutations) that are not mentioned in the literature about PC or BPH and are unique to our BPH sample cohort. When viewing each mutation seperatly, most of th cases only one occurrence per mutation had heteroplasmy higher than 5%, and in many samples, on average, it was 1% high. Altogether, SNPs were found: 2 in ND1, 2 in ND2, 2 in tRNA-Ala, 2 in CO1, 3 CO2, 1 in CO3, 1 in ND4L, 4 in ND4, 2 in ND5, 2 in ND5, 2 in CoQ genes and 3 in 12S, 4 in 16S and 1 in HVS1 regions.

**Table 2.** Variants that are unique in our sample cohort.

| POS | Allele change | Sample count that had the mutation (occurrences) | Heteroplasmy, homoplasmy (cases) |  | Gene | Amino acid change | Type of amino acid change |
| --- | --- | --- | --- | --- | --- | --- | --- |
|  |  |  | Blood | Prostate tissue |  |  |  |
| 1104 | A>G | 2 | 5%<br>1% | no<br>no | 12S | rRNA | - |
| 1219 | T>C | 3 | 1%<br>7%<br>no | no<br>no<br>2% | 12S | rRNA | - |
| 1260 | A>G | 2 | 30%<br>no | no<br>2% | 12S | rRNA | - |
| 1849 | C>T | 1 | 45% | 80% | 16S | rRNA | - |
| 2105 | G>A | 1 | no | 10% | 16S | rRNA | - |
| 2729 | T>C | 4 | 100%<br>1%<br>1%<br>no | 100%<br>no<br>no<br>1% | 16S | rRNA | - |
| 2931 | A>G | 4 | 97%<br>1%<br>1%<br>1% | 100%<br>no<br>no<br>no | 16S | rRNA | - |
| 3620 | T>C | 2 | 8%<br>no | 1%<br>1% | ND1 | I105T | nonpolar → polar |
| 3914 | G>A | 2 | no<br>no | 13%<br>1% | ND1 | G203E | nonpolar → acidic |
| 4986 | A>G | 3 | no<br>no<br>1% | 1%<br>6%<br>no | ND2 | T173A | polar → nonpolar |
| 5033 | A>G | 4 | no<br>no<br>98%<br>1% | 1%<br>1%<br>100%<br>no | ND2 | - | - |
| 5609 | T>C | 7 | no<br>no<br>no<br>no<br>1%<br>no<br>1% | 1%<br>14%<br>1%<br>1%<br>no<br>1%<br>no | tRNA-Ala | T-stem | - |
| 5610 | G>A | 6 | no<br>no<br>1%<br>no<br>1%<br>no | 1%<br>6%<br>no<br>1%<br>no<br>1% | tRNA-Ala | T-stem | - |
| 6043 | T>C | 4 | 1%<br>no<br>no<br>1% | no<br>7%<br>1%<br>no | CO1 | L47P | nonpolar → nonpolar |
| 7179 | T>C | 6 | N/A<br>no<br>5%<br>no<br>1%<br>no | 1%<br>1%<br>no<br>1%<br>no<br>1% | CO1 | F426L | nonpolar → nonpolar |
| 7647 | T>C | 1 | 1% | 11% | CO2 | I21T | nonpolar → polar |
| 7874 | A>G | 2 | N/A<br>no | 24%<br>1 | CO2 | I97V | nonpolar → nonpolar |
| 7923 | A>G | 1 | 100% | 99% | CO2 | Y113C | polar → polar |
| 8214 | T>C | 4 | no<br>1%<br>1%<br>5% | 1%<br>no<br>no<br>1% | CO2 | V210A | nonpolar → nonpolar |
| 9500 | C>T | 3 | 2%<br>1%<br>100% | no<br>no<br>100% | CO3 | - | - |
| 10572 | G>A | 2 | no<br>no | 1%<br>34% | ND4L | G35Stop | nonpolar → stop |
| 10791 | T>C | 2 | no<br>no | 1%<br>8% | ND4 | L11S | nonpolar → polar |
| 11311 | A>G | 1 | 100% | 100% | ND4 | - | - |
| 11814 | T>C | 3 | 15%<br>no<br>1% | no<br>3%<br>no | ND4 | L352P | nonpolar → nonpolar |
| 12015 | T>C | 3 | 14%<br>1%<br>5% | 3%<br>no<br>no | ND4 | L419P | nonpolar → nonpolar |
| 14093 | T>C | 2 | 6%<br>2% | no<br>no | ND5 | L586P | nonpolar → nonpolar |
| 14126 | T>C | 4 | no<br>1%<br>2%<br>no | 10%<br>no<br>no<br>3% | ND5 | L597P | nonpolar → nonpolar |
| 14995 | C>T | 2 | 100%<br>1% | 100%<br>no | CoQ | - | - |
| 15164 | T>C | 1 | 100% | 100% | CoQ | F140L | nonpolar → nonpolar |
| 16128 | C>T | 4 | 1%<br>10%<br>1%<br>1% | no<br>2%<br>no<br>no | D-loop,<br>HVS1 | noncoding | - |

\*These SNPs are not associated with any of a mitochondrial haplogroup by PhyloTree.org.
\*\* If heteroplasmy level for a SNP was detected above 5% threshold, then for those SNPs the threshold was removed to get more precise analysis.

Only in 15 (16%) individual occurrences of 95 (32 different mutations (**Table 1 plus 2**)) were detected in both tissue types. In 2 patients, data for blood tissue were not available. For 41 (43%) of those mutation occurrences, it was detected in prostate tissue, but not blood; and 39 (41%) the occurrences were detected in blood cells but not prostate tissue (P=0.9512) (**Supplementary Figure 1**). The difference in heteroplasmy level in 15 mutation occurrences between prostate tissue and blood samples was from 0-35%. On average, the heteroplasmy level was 32% in prostate tissue and 28% in blood cells (P=0.95). For 6 occurrences, heteroplasmy was higher in blood tissue, but for 4 in prostate tissue; in 5 occurrences, it was homoplasmy (P=0.6703, Chi-square=0.8000, df=2). These mutations were considered unique for our BPH samples.

**Supplementary Table 1** represents 63 mtDNA SNPs in PBH samples that have been associated with PC or BPH in previous publications (reviewed in Kalsbeek et al., 2017) and were also detected in our sample cohort but are also associated with mtDNA haplogroups by HaploGrep 2.0; thus, these were not considered as unique SNPs for our BPH patients. Likewise, 51 SNPs that were detected in the BPH group but are associated with another mtDNA haplogroup (not specific to a sample’s haplogroup) were not considered unique (**Supplementary Table 2**). Also, 89 mutations which are not associated with the haplogroups, but were detected in the samples of the general Latvian population, were also not considered as unique mutations for BPH patients (**Supplementary Table 3**). Individuals from the general Latvian population did not have cancer or benign tumours at the time of DNA material collection but were, on average, 24 years younger than the BPH group. That might mean that some of the variants from this Supplementary table 3 might biasedly be misplaced as some male individuals might develop diseases later in life. However, this was the data from the general population available for the study.

## Discussion

Unfortunately, BPH is not a completely understood disorder despite the fact that it affected 66.9% world’s population of men as of 2017 (Launer et al., 2021). And there is a lack of studying BPH in connection with cell ageing hallmarks in one sample cohort. Our findings show that there are changes in TL and mtDNA for BPH patients, and unique variants of mtDNA can be found among BPH patients.

We showed that BPH prostate tissue had the longest telomeres, which might explain why apoptosis and cell death are not initiated. Hyperplasia is caused when cell count increases in the organ, which should cause shorter telomeres in BPH (as cells divide more frequently) as in the literature described (Cheng et al., 2020; Sommerfeld et al., 1996). However, in our and another study, BPH had the longest telomeres compared with normal or cancerous tissue (Heaphy et al., 2010). In this case, telomerase might be involved in telomere prolongation. Although we did not measure telomerase activity and in literature, there are very controversial results showing both increased activity and no activity at all (Kamradt et al., 2003; Rane et al., 2016; Sommerfeld et al., 1996; Wymenga et al., 2000; Zhang et al. 1998; Mahjoub and Krams 2006). In our BPH group, we had a high heterogeneity which might also suggest another hypothesis that there are telomere circles in benign tissue. Telomere circles have been found in healthy human tissue with the Circle-seq method by which extrachromosomal circular DNA (eccDNA) is found (Møller et al., 2018). eccDNA can be especially abounded in diseased tissue because of genomic instability (Kumar et al., 2017). The potential telomere circles in BPH tissue can be detected by qPCR as a stronger signal. Another possibility is that telomeres are prolonged by the alternative lengthening of telomeres (ATL) (Bower et al., 2012; Cesare and Griffith, 2004), although it is unlikely. Even though ATL has been reported in normal mammalian somatic cells (Neumann et al., 2013; reviewed in Zhdanova and Rubtsov, 2016). Also, the blood cells of our BPH group showed disturbed TL regulation as it had more heterogeneous and longer telomeres than the blood cells of the healthy group. In the literature, PC patients with long leukocyte telomeres had poorer survival (Svenson et al., 2017). The only PC reference sample had shorter telomeres than BPH prostate tissue, which is as described in the literature; although we had only one sample, it might show that measurements were accurate. It seems like prolonged telomeres might be one of the factors that are involved in the development of BPH and potentially can be used as a biomarker in combination with other markers.

Some researchers have found mutations in *TP53* in BPH (Meyers et al., 1993; Schlechte et al., 1998; Wertz et al., 1996). We found the 7676154C>G (rs1042522) mutation in the 4^th^ exon. This mutation has been previously associated with some cancers (Chen et al., 2015; Huang et al., 2019) but not with PC (Fan et al., 2017). Also, our study showed that this mutation is inherited and is in both BPH and healthy groups in a similar frequency, which proves this is not a risk factor for the development of BPH. This agrees with the literature, where 23 genome-wide significant variants are associated with BPH and non in *TP53* (Gudmundsson et al., 2018).

Not many studies have been conducted regarding mtDNA amount and BPH. One of the previous reports showed reduced mtDNA amount in BPH patients and blood cells compared to normal tissues in African American and Caucasian-American men (Koochekpour et al., 2013). In addition, BPH patients had slightly less mtDNA amount than PC patients (Abhishek et al., 2017). In our study, BPH blood cells had more mtDNA than controls’ blood, and BPH prostate tissue had the highest amount. The one PC reference sample showed a higher amount than BPH prostate tissue which is in agreement with the literature (Abhishek et al., 2017; Zhou et al., 2014). Increased mtDNA amount might promote cell survival by accelerating cell proliferation and inhibiting apoptosis, as in microsatellite stable colorectal cancer (MSS CRC) cells (Sun et al., 2018). This might explain why in BPH prostate tissue, mtDNA amount is as high, thus allowing cells to proliferate at an abnormal rate. In some cases, increased mtDNA content correlated with a protective effect, for example, in glioma patients (Zhang et al., 2015). This might suggest why these BPH cells are not becoming highly aggressive and cancerous. All the contradicting studies on mtDNA amount and different cancers (Hu et al., 2016) and diseases (e.g., (Yue et al., 2018)) make it hard to conclude the exact role of it in the development of BPH, but it is changed in BPH patients.

Regarding mtDNA haplogroup U, BPH had no statistically significant association with it in our sample cohort. Although, in our BPH group, there was 10% more of haplogroup U than in the Latvian population. More individuals with haplogroup U have been seen in other studies regarding PC than healthy individuals (Booker et al., 2006; McCrow et al., 2016). To prove any association, a bigger sample size cohort of BPH should be tested to make concrete conclusions.

Two variants in Table 1, G3047A and G5590A might be involved in BPH development as well as PC development later in life, but more samples and follow up must be done. These SNPs have been previously associated with PC (Ju et al., 2014). Although if the heteroplasmy level is low, that might not be enough to influence the function of mitochondria (Saneto, 2017), and in our cohort, only one or two samples had them higher than 5%.

Many amino acid changes (Table 2) can heavily interfere with protein 3D structure and function (Lloyd and McGeehan, 2013), including changes in 12S and 16S, causing different diseases (Elson et al., 2015; Smith et al., 2014). But most of the mutations we found are in a heteroplasmic state and at a very low level. Only a few homoplasmic mutations, or the ones with enough high heteroplasmy levels, like A1260G and C1849T, might be high enough to cause a disease (reviewed in Rossignol et al., 2003). In our sample cohort, it seems that heteroplasmy is very dynamic between the tissues (Table 2), but it does not show a higher level or more frequent heteroplasmy in prostate tissue in comparison with blood. This does not mean that heteroplasmy in prostate tissue could not cause BPH if it is not in the blood cells, but probably enough high level should be present. There is no research describing the heteroplasmy level in BPH so far in our knowledge, which means there is a necessity for more of these studies. More functional studies or research with more samples should be conducted to prove a specific connection between our found mutations and the development of BPH.

Many of the mutations in our cohort that have been associated with PC or other diseases (McCrow et al., 2016; McGeehan et al., 2018; reviewed in Kalsbeek et al., 2017; Lott et al., 2013) were also associated with one of the mtDNA haplogroups but not to the specific haplogroup of the sample. Although some mutations can change the protein function or mtDNA regulation (Supplementary Table 1, 2, 3), we did not look into these as they are not specific to our BPH patients’ group.

In conclusion, our BPH blood samples had prolonged telomeres in comparison to healthy blood samples, and BPH prostate tissue had the longest telomeres; thus, TL might be involved in the development of BPH. Similarly, changes in mtDNA amount might contribute in some way as we saw a higher amount in BPH patients’ blood cells than in healthy individuals, and BPH prostate tissue had the highest. Although, it is not clear if it has a negative or compensatory effect as studies about mtDNA amount are very controversial. Regarding the mtDNA genome, some of the heteroplasmic or homoplasmic variations might contribute to the development of BPH as they cause hazardous changes in proteins or regulatory regions of mtDNA maintenance. Few of them also have enough high heteroplasmy levels that might affect mitochondrial functions, and some have been associated with PC previously. We did not find any changes in the *TP53* gene that could be causing the BPH. The limitations of the study were the small number of tissue samples from BPH patients, the younger average age and the relatively small sample amount for the group of the general Latvian population. Although none of the control samples had been diagnosed with cancer at the time of DNA sampling, they might develop the diseases later in life. As there are not many studies addressing the issues in BPH, this study provides insights into the factors viewed and provides a platform for future studies to prove any of the involvements and interactions in the development of BPH.

## Acknowledgement

We thank SIA GenEra and the staff for technical support regarding qPCR assays. Also, administrators of the Genome Database of the Latvian Population for providing DNA samples for controls used in this study.

---

**Supplementary figure 1.**
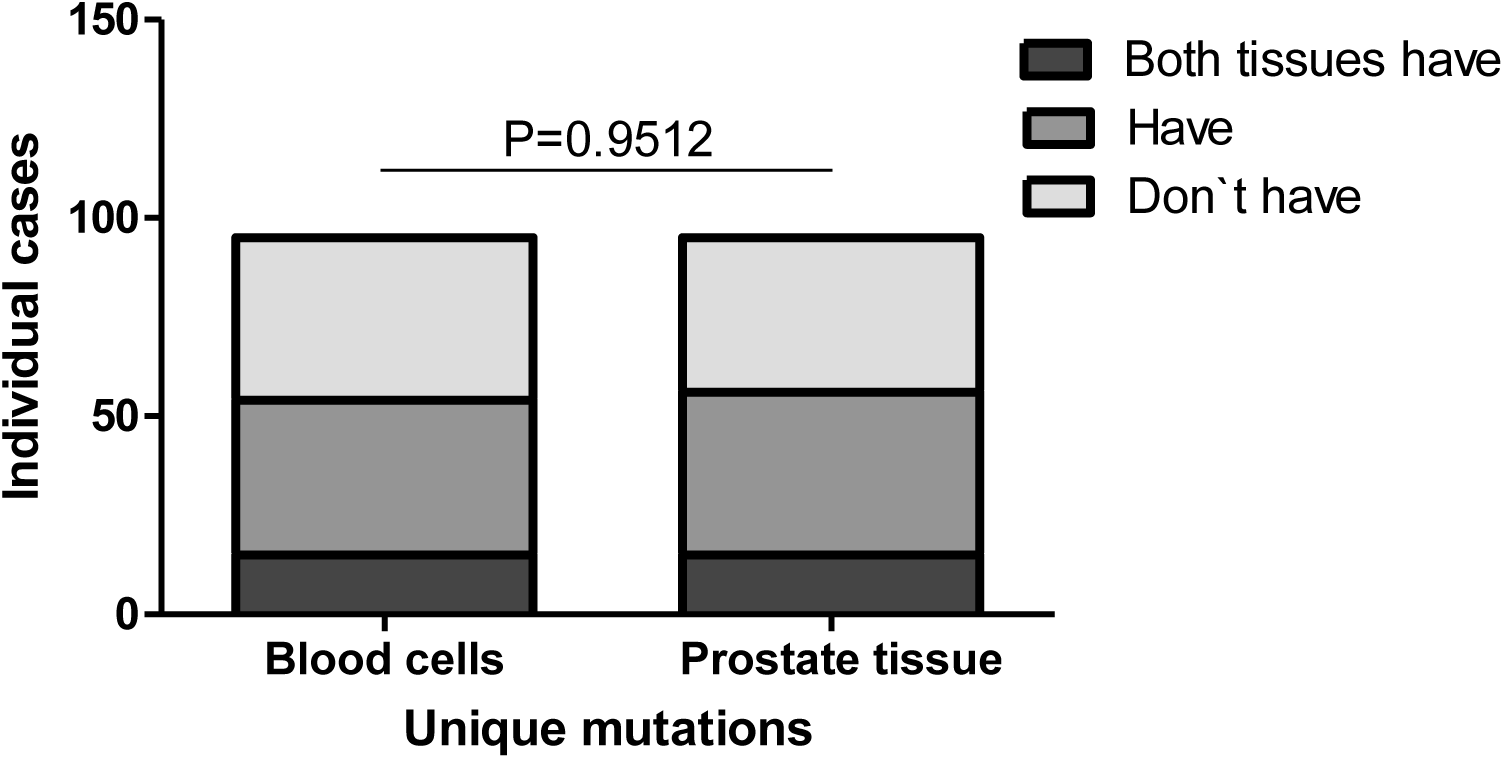
Comparison of frequency of heteroplasmy in BPH patients. Critical Chi-square value=0.1, df=2, α=0.05.

**Supplementary figure 2.**
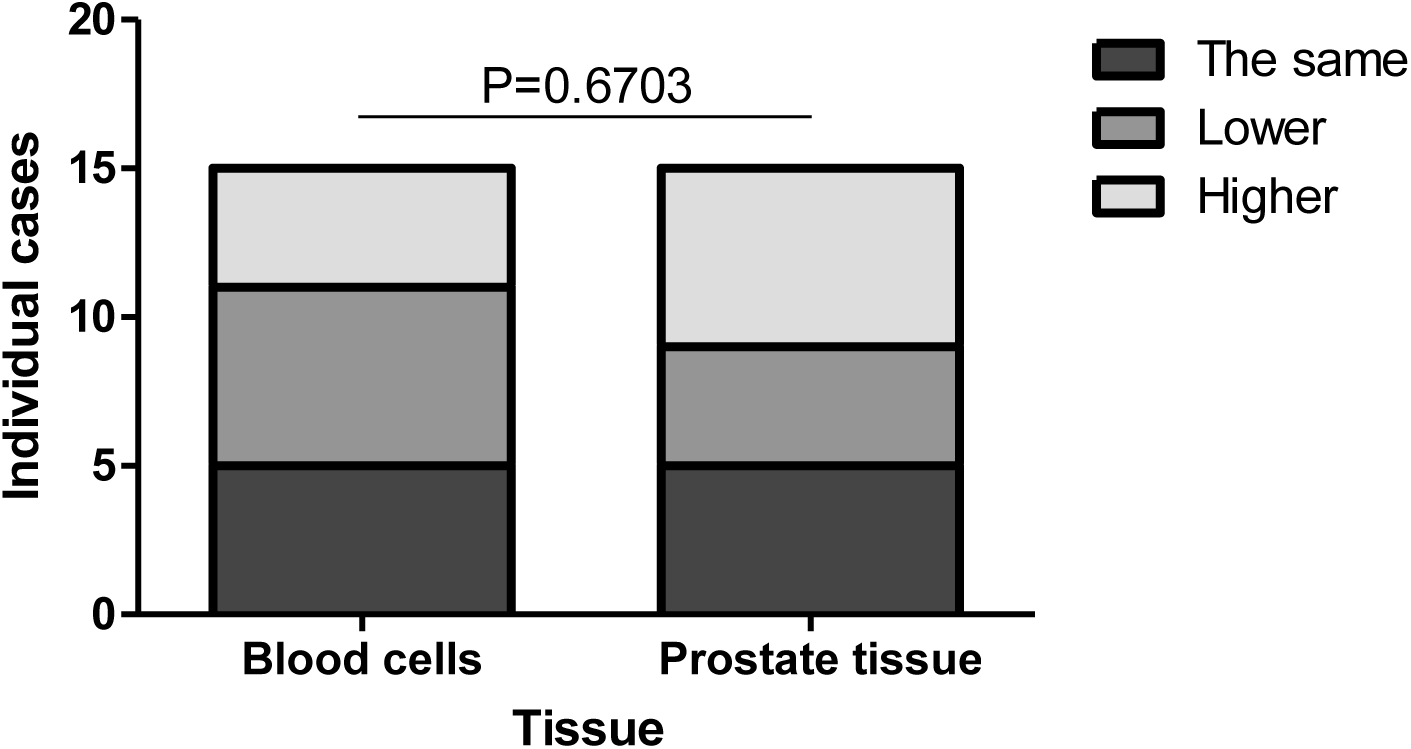
Comparison of level of heteroplasmy in the 15 individual cases. Critical Chi-square value=0.8, df=2, α=0.05.

**Supplementary Table 1.**
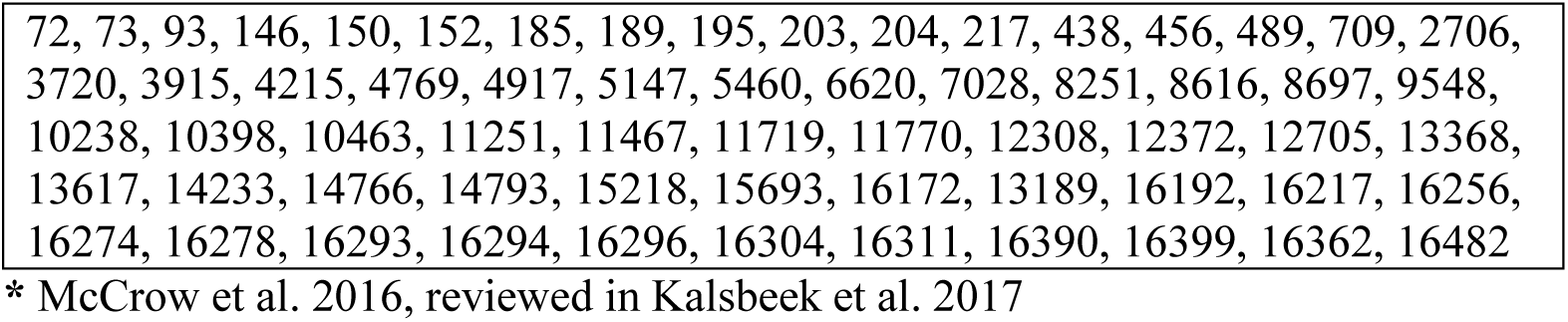
SNPs’ positions in mtDNA that were detected in BPH samples, were previously associated with prostate cancer*, and are associated with the specific samplès mtDNA haplogroup.

**Supplementary Table 2.** SNPs that are associated with mtDNA haplogroups but do not belong to the specific BPH samplès mtDNA haplogroup.

| SNPs that are: • in BPH samples<br>• not in the publications*<br>• not in the control group of a general population<br>• associated with mtDNA haplogroups | SNPs that are: • in BPH samples<br>• in the publications*<br>• not in the control group of a general population<br>• associated with mtDNA haplogroups |
| --- | --- |
| Homoplasmy:<br>A153G, T246C, C296T, T2352C, C2639T, T3618C, C4350T, C5461T, T6221C, A6359G, G7830A (R82H), T7837C, T8167C, A8291G, C8778T, G8865A, G8994A, G9196A (D22AN), T10454C, T10885C, T11485C, C11536T, C11665T, G12007A, A12397G, T12441C, G12618A, T13443C, C14115T G15257A, T15889C, T16086C, T16092C, T16209C | Homoplasmy:<br>G7912A, G8723A |
| Heteroplasmy:<br>C151T, C3921A, A4257G, A4435G, T4977C, C5360T, G7337A, A10598G, T11935C, G13980A, T14180C, T15601C, T16124C | Heteroplasmy:<br>G1415A, T16093C |

\* McCrow et al. 2016, reviewed in Kalsbeek et al. 2017

**Supplementary Table 3.** Other SNPs in BPH samples detected.

| POS | Allele change | Variant frequency |  |  |  | Heteroplasmy level (%)<br>Average (min-max) |  |  |  | Gene | Amino acid change | Type of amino acid change | Previously reported different changes of the position | Article | Predicti on in the publicat ion |
| --- | --- | --- | --- | --- | --- | --- | --- | --- | --- | --- | --- | --- | --- | --- | --- |
|  |  | Control (n=50) | Blood (n=30) | Prostate (n=32) | P value | Control (n=50) | Blood (n=30) | Prostate (n=32) | P value |  |  |  |  |  |  |
| 65 | TG>T | 0.72 | 0.73 | 0.56 | 0.2488 | 3 (1-10) | 4 (1-27) | 5 (2-18) | 0.194 | D-loop, HVSII | noncoding | - | - | Ju et al. 2014 | Primary PC |
| 65 | T>TG | 0.94 | 0.93 | 0.91 | 0.8398 | 9 (2-27) | 6 (1-12) | 7 (2-16) | 0.0676 | D-loop, HVSII | noncoding | - | - | - | - |
| 285 | C>CA | 0.62 | 0.72 | 0.56 | 0.4145 | 6 (2-20) | 3 (1-11) | 6 (2-13) | 0.0027 | D-loop, HVSII | noncoding | - | - | - | - |
| 302 | AC>A | 0.36 | 0.59 | 0.16 | 0.0022 | 7 (2-15) | 6 (2-13) | 10 (3-20) | 0.2839 | D-loop, HVSII | noncoding | - | - | - | - |
| 308 | C>C<sub>n</sub> | 0.83 | 0.83 | 0.50 | 0.0022 | 17 (2-67) | 11 (2-29) | 10 (3-25) | 0.0226 | D-loop, HVSII | noncoding | - | C>T, het | Kalsbeek et al. 2016 | Primary PC |
| 309 | C>C<sub>n</sub> | 0.88 | 1.00 | 0.81 | 0.0573 | 41 (2-94) | 38 (6-100) | 41 (6-92) | 0.9218 | D-loop, HVSII | noncoding | - | - | Kloss-Brandstätter et al. 2010 | Primary PC |
| 309 | C>T | 0.37 | 0.48 | 0.38 | 0.5750 | 6 (2-13) | 5 (2-7) | 12 (2-79) | 0.2339 | D-loop, HVSII | noncoding | - | - | - | - |
| 310 | T>C | 0.98 | 1.00 | 0.91 | 0.1148 | 57 (6-100) | 51 (15-100) | 43 (7-100) | 0.1624 | D-loop, HVSII | noncoding | - | - | Gómez-Zaera et al. 2006 | Primary PC |
| 310 | T>TC | 0.82 | 0.90 | 0.97 | 0.1328 | 47 (2-100) | 58 (3-92) | 53 (4-100) | 0.3323 | D-loop, HVSII | noncoding | - | - | Gómez-Zaera et al. 2006 | Primary PC |
| 316 | G>C | 0.13 | 0.27 | 0.09 | 0.1434 | 2<br>(0.1-4) | 3<br>(1-6) | 6<br>(2-11) | 0.1178 | D-loop | noncoding | - | - | Kalsbeek et al. 2016<br>Kalsbeek et al. 2017 | Primary PC bone metastasis |
| 316 | G>A | 0.05 | 0.03 | 0.03 | 0.8916 | 6<br>(4-8) | 94<br>(94-94) | 89<br>(89-89) | 0.1940 | D-loop, HVSII | nancoding | - | - | - | - |
| 451 | A>AT | 0.03 | 0.03 | 0.06 | 0.8170 | 92<br>(92-92) | 84<br>(84-84) | 96<br>(93-100) | - | D-loop, HVSIII | noncoding | - | - | - | - |
| 513 | GCA>G | 0.31 | 0.27 | 0.19 | 0.5098 | 77<br>(3-100) | 30<br>(3-100) | 33<br>(4-89) | 0.0393 | D-loop, HVSIII | noncoding | - | - | - | - |
| 513 | G>GCA | 0.05 | 0.07 | 0.06 | 0.9341 | 38<br>(14-62) | 72<br>(68-75) | 54<br>(40-68) | 0.4449 | D-loop, HVSIII | noncoding | - | - | Ju et al. 2014 | Primary PC |
| 567 | A>AC<sub>n</sub> | 0.46 | 0.33 | 0.41 | 0.5502 | 9<br>(1-83) | 16<br>(2-57) | 21<br>(3-77) | 0.3342 | D-loop, HVSIII | noncoding | - | - | Kalsbeek et al. 2017 | bone metastasis |
| 955 | AC>C | 0.20 | 0.28 | 0.09 | 0.1858 | 1<br>(0.1-4) | 1<br>(0.1-2) | 4<br>(3-7) | 0.0047 | 12S | rRNA | - | - | Ju et al. 2014 | Primary PC |
| 955 | A>AC<sub>n</sub> | 0.08 | 0.07 | 0.06 | 0.9399 | 3<br>(2-4) | 70<br>(50-89) | 70<br>(46-94) | 0.0130 | 12S | rRNA | - | - | - | - |
| 992 | T>TA | 0.65 | 0.50 | 0.69 | 0.2615 | 3<br>(1-8) | 2<br>(1-5) | 3<br>(1-6) | 0.0017 | 12S | rRNA | - | T>C, het, rRNA | Ju et al. 2014 | Primary PC |
| 1369 | T>C | 0.04 | 0 | 0.03 | 0.5529 | 8<br>(1-15) | 0 | 15<br>(15-15) | - | 12S | rRNA | - | - | - | - |
| 2129 | G>GA | 0.58 | 0.63 | 0.66 | 0.7667 | 4<br>(1-11) | 2<br>(1-5) | 3<br>(1-10) | 0.0036 | 16S | rRNA | - | - | - | - |
| 2150 | T>del | 0.16 | 0.41 | 0.13 | 0.0111 | 2<br>(0.1-5) | 2<br>(1-5) | 2<br>(1-3) | 0.5281 | 16S | rRNA | - | - | Kloss-Brandstätter et al. 2010 | Primary PC |
| 2150 | T>TA | 0.68 | 0.87 | 0.69 | 0.1506 | 7<br>(1-16) | 4<br>(1-9) | 5<br>(2-12) | 0.0007 | 16S | rRNA | - | - | - | - |
| 2226 | TA>T | 0.50 | 0.53 | 0.25 | 0.0391 | 5<br>(1-29) | 3<br>(1-7) | 3<br>(1-7) | 0.2348 | 16S | rRNA | - | - | - | - |
| 2397 | C>A | 0.04 | 0.03 | 0.03 | 0.9752 | 50<br>(0.1-100) | 100<br>(100-100) | 100<br>(100-100) | - | 16S | rRNA | - | - | - | - |
| 2456 | TA>T | 0.71 | 0.86 | 0.72 | 0.2922 | 4<br>(1-11) | 4<br>(1-9) | 4<br>(1-9) | 0.5068 | 16S | rRNA | - | - | - | - |
| 2456 | T>TA | 0.90 | 0.93 | 0.88 | 0.7413 | 7<br>(3-23) | 6<br>(2-11) | 7<br>(2-19) | 0.7634 | 16S | rRNA | - | - | - | - |
| 3380 | G>GA | 0.54 | 0.67 | 0.63 | 0.5002 | 2<br>(0.1-8) | 2<br>(1-6) | 2<br>(1-3) | 0.6313 | ND1 | frameshift | - | G>A, het,<br>R25Q | Ju et al. 2014 | Primary<br>PC |
| 3565 | A>AC | 0.50 | 0.27 | 0.44 | 0.1740 | 2<br>(0.1-5) | 2<br>(2-3) | 4<br>(1-11) | 0.1045 | ND1 | frameshift | - | - | Ju et al. 2014 | Primary<br>PC |
| 3565 | AC>A | 0.63 | 0.50 | 0.53 | 0.5468 | 3<br>(1-9) | 3<br>(1-7) | 3<br>(1-6) | 0.8101 | ND1 | frameshift | - | - | - | - |
| 3912 | A>AG | 0.68 | 0.77 | 0.75 | 0.6492 | 4<br>(1-8) | 3<br>(1-8) | 3<br>(1-7) | 0.008 | ND1 | frameshift | - | - | - | - |
| 4175 | G>GA | 0.88 | 0.87 | 0.91 | 0.8786 | 6<br>(1-14) | 3<br>(1-9) | 2<br>(1-5) | < 0.0001 | ND1 | frameshift | - | - | - | - |
| 4393 | C>A | 0.06 | 0.03 | 0.06 | 0.8464 | 34<br>(0.1-100) | 100<br>(100-100) | 51<br>(1-100) | - | tRNA-Gln | tRNA | - | - | - | - |
| 4547 | AT>A | 0.48 | 0.80 | 0.81 | 0.0014 | 2<br>(1-4) | 2<br>(1-7) | 2<br>(1-7) | 0.2179 | ND2 | frameshift | - | - | - | - |
| 4547 | A>AT | 0.58 | 0.63 | 0.69 | 0.6144 | 3<br>(1-6) | 2<br>(1-3) | 2<br>(1-5) | 0.0930 | ND2 | frameshift | - | - | - | - |
| 4604 | CA>C | 0.67 | 0.90 | 0.84 | 0.0316 | 3<br>(1-7) | 3<br>(1-8) | 3<br>(1-8) | 0.6465 | ND2 | frameshift | - | - | - | - |
| 4604 | C>CA | 0.90 | 0.97 | 0.97 | 0.3090 | 6<br>(1-16) | 5<br>(1-9) | 5<br>(1-11) | 0.4624 | ND2 | frameshift | - | - | - | - |
| 4855 | T>C | 0.16 | 0.07 | 0.06 | 0.2670 | 1<br>(0.1-1) | 1<br>(1-1) | 4<br>(1-6) | 0.0525 | ND2 | L129P | nonpolar →<br>nonpolar | - | - | - |
| 4867 | GA>G | 0.49 | 0.90 | 0.91 | <<br>0.0001 | 3<br>(0.1-10) | 5<br>(1-11) | 5<br>(1-15) | 0.0102 | ND2 | frameshift | - | - | - | - |
| 5231 | G>GC | 0.55 | 0.57 | 0.75 | 0.1612 | 3<br>(1-9) | 3<br>(1-7) | 2<br>(1-4) | 0.3752 | ND2 | frameshift | - | - | - | - |
| 5281 | C>CA | 0.71 | 0.75 | 0.81 | 0.6119 | 3<br>(1-9) | 3<br>(1-6) | 3<br>(1-8) | 0.4478 | ND2 | frameshift | - | - | - | - |
| 5823 | AG>A | 0.68 | 0.80 | 0.97 | 0.0068 | 4<br>(1-14) | 4<br>(1-8) | 4<br>(1-9) | 0.8846 | tRNA-Cys | Acceptor<br>stem | - | - | - | - |
| 5894 | AC>A | 0.14 | 0.27 | 0.19 | 0.3726 | 2<br>(0.1-6) | 1<br>(1-2) | 2<br>(1-3) | 0.2652 | noncoding | noncoding | - | - | Ju et al. 2014 | Primary<br>PC |
| 5894 | A>AC | 0.56 | 0.40 | 0.53 | 0.3873 | 2<br>(0.1-5) | 9<br>(1-95) | 7<br>(1-93) | 0.4431 | noncoding | noncoding | - | - | Ju et al. 2014 | Primary<br>PC |
| 6691 | GA>G | 0.66 | 0.80 | 0.94 | 0.0123 | 3<br>(1-15) | 2<br>(1-4) | 2<br>(1-5) | 0.1742 | CO1 | frameshift | - | N/A | Parr et al.<br>2006 | Primary<br>PC |
| 6691 | G>GA | 0.98 | 1.00 | 1.00 | - | 11<br>(1-30) | 8<br>(5-15) | 7<br>(2-14) | < 0.0001 | CO1 | frameshift | - | - | - | - |
| 7396 | GC>G | 0.95 | 0.97 | 1.00 | 0.4633 | 12<br>(2-26) | 12<br>(3-25) | 10<br>(2-25) | 0.3852 | CO1 | frameshift | - | - | - | - |
| 7396 | G>GC | 0.66 | 0.90 | 0.72 | 0.0621 | 5<br>(2-14) | 6<br>(2-14) | 5<br>(1-10) | 0.0765 | CO1 | frameshift | - | - | - | - |
| 7504 | A>AT | 0.56 | 0.87 | 0.78 | 0.0139 | 4<br>(1-14) | 4<br>(1-9) | 5<br>(1-10) | 0.2574 | tRNA-SerUCN | DHU-stem | - | - | - | - |
| 8016 | T>TC | 0.62 | 0.87 | 0.88 | 0.0110 | 3<br>(1-7) | 2<br>(1-8) | 2<br>(1-6) | 0.5808 | CO2 | frameshift | - | - | - | - |
| 8232 | TA>T | 0.82 | 0.97 | 0.97 | 0.0384 | 10<br>(1-63) | 8<br>(2-15) | 7<br>(1-16) | 0.1474 | CO2 | frameshift | - | - | - | - |
| 8496 | T>TA | 0.44 | 0.37 | 0.31 | 0.5840 | 3<br>(1-9) | 4<br>(1-8) | 5<br>(1-9) | 0.0715 | ATP8 | frameshift | - | - | - | - |
| 9432 | C>CA | 0.44 | 0.40 | 0.72 | 0.0179 | 2<br>(0.1-4) | 2<br>(1-7) | 2<br>(1-8) | 0.9791 | CO3 | frameshift | - | - | - | - |
| 9474 | G>GA | 0.12 | 0.27 | 0.25 | 0.1855 | 8<br>(5-12) | 6<br>(1-7) | 7<br>(4-12) | 0.2693 | CO3 | frameshift | - | G>A, het,<br>E90K | Lindberg et<br>al. 2013 | Primary<br>PC |
| 9476 | AG>A | 0.06 | 0.17 | 0.19 | 0.1694 | 3<br>(3-4) | 2<br>(1-4) | 2<br>(1-6) | 0.3607 | CO3 | frameshift | - | - | - | - |
| 9477 | G>GT | 0.66 | 0.77 | 0.75 | 0.5155 | 7<br>(1-20) | 7<br>(4-12) | 6<br>(1-13) | 0.6286 | CO3 | frameshift | - | G>C, het,<br>V91L | Arnold et al.<br>2015 | soft<br>tissue<br>metastasis |
| 9494 | A>AT | 0.67 | 0.33 | 0.44 | 0.0121 | 2<br>(0.1-6) | 2<br>(1-7) | 2<br>(1-5) | 0.5914 | CO3 | frameshift | - | - | - | - |
| 9768 | AT>A | 1.00 | 1.00 | 1.00 | - | 23<br>(5-67) | 22<br>(3-38) | 23<br>(5-60) | 0.7948 | CO3 | frameshift | - | - | - | - |
| 9794 | A>AT | 0.67 | 0.77 | 0.78 | 0.4624 | 4<br>(1-22) | 4<br>(1-17) | 3<br>(1-7) | 0.5209 | CO3 | frameshift | - | - | - | - |
| 9905 | TG>T | 0.62 | 0.93 | 0.88 | 0.0014 | 10<br>(3-36) | 7<br>(2-18) | 7<br>(3-17) | 0.2185 | CO3 | frameshift | - | - | - | - |
| 10348 | TC>T | 0.94 | 1.00 | 1.00 | - | 12<br>(1-33) | 13<br>(6-26) | 11<br>(4-22) | 0.4700 | ND3 | frameshift | - | - | - | - |
| 10813 | CA>C | 0.66 | 0.80 | 0.66 | 0.3534 | 1<br>(0.1-4) | 1<br>(0.1-3) | 1<br>(0.1-2) | 0.1634 | ND4 | frameshift | - | - | Ju et al. 2014 | Primary<br>PC |
| 10813 | C>CA | 0.98 | 0.93 | 0.97 | 0.5456 | 4<br>(1-14) | 2<br>(0.1-5) | 2<br>(1-5) | < 0.0001 | ND4 | frameshift | - | - | - | - |
| 10879 | AT>A | 0.63 | 0.72 | 0.47 | 0.1195 | 3<br>(1-8) | 4<br>(1-11) | 7<br>(1-16) | 0.0087 | ND4 | frameshift | - | - | - | - |
| 10879 | A>AT | 0.78 | 0.69 | 0.38 | 0.0015 | 5<br>(2-14) | 5<br>(2-11) | 5<br>(1-14) | 0.9673 | ND4 | frameshift | - | - | - | - |
| 10946 | A>AC | 0.67 | 0.69 | 0.31 | 0.0037 | 5<br>(1-12) | 5<br>(2-10) | 6<br>(3-10) | 0.7727 | ND4 | frameshift | - | - | - | - |
| 11031 | G>GA | 0.93 | 1.00 | 0.88 | 0.1455 | 14<br>(3-45) | 13<br>(2-28) | 10<br>(2-50) | 0.1875 | ND4 | frameshift | - | - | - | - |
| 11093 | G>C | 0.56 | 0.55 | 0.50 | 0.8702 | 5<br>(1-16) | 3<br>(1-6) | 5<br>(1-13) | 0.0950 | ND4 | A112P | nonpolar →<br>nonpolar | - | - | - |
| 11711 | G>A | 0.38 | 0.57 | 0.69 | 0.0204 | 2<br>(0.1-6) | 1<br>(1-4) | 3<br>(1-6) | 0.0468 | ND4 | A318T | nonpolar →<br>polar | - | Ju et al. 2014 | Primary<br>PC |
| 11826 | C>CT | 0.50 | 0.59 | 0.59 | 0.6358 | 3<br>(0.1-9) | 2<br>(1-4) | 3<br>(1-9) | 0.0286 | ND4 | frameshift | - | - | - | - |
| 11866 | A>AC | 0.83 | 0.83 | 0.66 | 0.1538 | 6<br>(1-18) | 3<br>(1-6) | 4<br>(1-11) | 0.0004 | ND4 | frameshift | - | - | - | - |
| 12384 | TC>T | 0.95 | 0.90 | 0.78 | 0.0992 | 12<br>(2-33) | 12<br>(3-47) | 12<br>(3-22) | 0.9898 | ND5 | frameshift | - | - | - | - |
| 12384 | T>TC | 0.44 | 0.69 | 0.25 | 0.0026 | 5<br>(2-13) | 6<br>(2-11) | 6<br>(2-9) | 0.8791 | ND5 | frameshift | - | - | - | - |
| 12417 | CA>C | 0.74 | 0.76 | 0.53 | 0.0906 | 9<br>(2-25) | 7<br>(2-14) | 8<br>(1-16) | 0.3123 | ND5 | frameshift | - | - | Ju et al. 2014 | Primary<br>PC |
| 12417 | C>CA | 1.00 | 0.97 | 1.00 | - | 36<br>(8-56) | 40<br>(19-54) | 34<br>(8-54) | 0.1123 | ND5 | frameshift | - | - | Ju et al. 2014 | Primary<br>PC |
| 13052 | GC>G | 0.64 | 0.72 | 0.59 | 0.5619 | 3<br>(1-15) | 3<br>(1-5) | 4<br>(1-9) | 0.2420 | ND5 | frameshift | - | - | - | - |
| 13230 | CA>C | 0.64 | 0.83 | 0.59 | 0.1193 | 3<br>(1-6) | 3<br>(1-7) | 4<br>(1-6) | 0.2855 | ND5 | frameshift | - | - | - | - |
| 13230 | C>CA | 0.89 | 1.00 | 0.88 | 0.1558 | 7<br>(1-19) | 6<br>(2-9) | 5<br>(2-10) | 0.0066 | ND5 | I298M | nonpolar →<br>nonpolar | - | - | - |
| 13299 | A>C | 0.69 | 0.83 | 1.00 | 0.0021 | 6<br>(1-13) | 12<br>(1-19) | 11<br>(4-22) | < 0.0001 | ND5 | Q321H | polar →<br>basic | - | - | - |
| 13304 | A>AG | 0.27 | 0.69 | 0.59 | 0.0005 | 4<br>(1-15) | 3<br>(1-6) | 3<br>(1-15) | 0.5302 | ND5 | frameshift | - | - | - | - |
| 13305 | C>G | 0.40 | 0.72 | 0.75 | 0.0015 | 2<br>(0.1-5) | 4<br>(1-8) | 4<br>(1-10) | 0.0274 | ND5 | H323Q | basic →<br>polar | - | - | - |
| 14355 | T>C | 0.26 | 0.40 | 0.41 | 0.3309 | 3<br>(1-6) | 5<br>(2-14) | 6<br>(2-10) | 0.0806 | ND6 | K107Q | basic →<br>polar | - | - | - |
| 14503 | T>TA | 0.85 | 0.93 | 0.84 | 0.5219 | 14<br>(4-30) | 15<br>(3-32) | 12<br>(3-25) | 0.1150 | ND6 | frameshift | - | - | - | - |
| 14750 | AC>A | 0.67 | 0.50 | 0.22 | 0.0013 | 4<br>(1-14) | 6<br>(2-16) | 8<br>(4-16) | 0.0429 | CoQ | frameshift | - | G>A, hom,<br>T2A | Arnold et al.<br>2015 | soft<br>tissue<br>metastasis |
| 14761 | CA>C | 0.79 | 0.83 | 0.75 | 0.7232 | 17<br>(3-50) | 15<br>(3-43) | 16<br>(2-40) | 0.8746 | CoQ | frameshift | - | - | - | - |
| 16182 | A>AC<sub>n</sub> | 0.08 | 0.07 | 0.09 | 0.9260 | 28<br>(8-50) | 13<br>(4-23) | 14<br>(1-36) | 0.6212 | D-loop,<br>HVSI | noncoding | - | A>C, het,<br>hom,<br>noncoding | Chen et al.<br>2002 | Primary<br>PC |
| 16183 | A>C | 0.16 | 0.10 | 0.09 | 0.6014 | 53<br>(2-100) | 35<br>(2-100) | 15<br>(1-36) | 0.4519 | D-loop,<br>HVSI | noncoding | - | A>G, hom,<br>noncoding | Chen et al.<br>2002,<br>Jerónimo et<br>al. 2001 | Primary<br>PC |
| 16183 | A>AC<sub>n</sub> | 0.21 | 0.07 | 0.06 | 0.0804 | 2<br>(1-4) | 2<br>(1-2) | 4<br>(3-5) | 0.0583 | D-loop,<br>HVSI | noncoding | - | - | Kalsbeek et<br>al. 2017,<br>Ju et al. 2014 | Primary<br>PC |
| 16183 | AC>A | 0.23 | 0.07 | 0.09 | 0.0866 | 10<br>(1-33) | 51<br>(48-53) | 30<br>(1-56) | 0.0064 | D-loop,<br>HVSI | noncoding | - | - | - | - |
| 16519 | T>C | 0.58 | 0.47 | 0.44 | 0.3910 | 96<br>(1-100) | 86<br>(1-100) | 93<br>(1-100) | 0.4544 | noncoding | noncoding | - | - | Kalsbeek et al.<br>2016,<br>Ju et al. 2014,<br>Chen et al. 2002,<br>Arnold et al. 2015 | bone<br>metastasis<br>Primary PC<br>Primary PC<br>Primary PC<br>soft tissue<br>metastasis |

These SNPs are not associated with any of a mitochondrial haplogroup by PhyloTree.org.
If heteroplasmy level for a SNP was detected above 5% threshold, then for those SNPs the threshold was removed to get more precise analysis.

